# A Framework of Multi-View Machine Learning for Biological Spectral Unmixing of Fluorophores with Overlapping Excitation and Emission Spectra

**DOI:** 10.1101/2024.08.07.607102

**Authors:** Ruogu Wang, Yunlong Feng, Alex M. Valm

## Abstract

The accuracy in assigning fluorophore identity and abundance, termed spectral unmixing, in biological fluorescence microscopy images remains challenging due to the unavoidable and significant overlap in emission spectra among fluorophores. In conventional laser scanning confocal spectral microscopy, fluorophore information is acquired by recording emission spectra with a single combination of discrete excitation wavelengths. As a matter of fact, organic fluorophores have not only unique emission spectral signatures but also have unique and characteristic excitation spectra. In this paper, we propose a generalized multi-view machine learning approach, which makes use of both excitation and emission spectra to greatly improve the accuracy in differentiating multiple highly overlapping fluorophores in a single image. By recording emission spectra of the same field with multiple combinations of excitation wavelengths, we obtain data representing these different views of the underlying fluorophore distribution in the sample. We then propose a framework of multi-view machine learning methods, which allows us to flexibly incorporate noise information and abundance constraints, to extract the spectral signatures of fluorophores from their reference images and to efficiently recover their corresponding abundances in unknown mixed images. Numerical experiments on simulated image data demonstrate the method’s efficacy in improving accuracy, allowing for the discrimination of 100 fluorophores with highly overlapping spectra. Furthermore, validation on images of mixtures of fluorescently labeled E. coli demonstrates the power of the proposed multi-view strategy in discriminating fluorophores with spectral overlap in real biological images.

## Introduction

Numerous biological systems exhibit intricate interactions among various subcomponents, and fluorescent labels are often employed to indicate the spatial distribution of these components within cells and tissues. Spectral imaging microscopes record fluorescence intensity data in discrete wavelength bands at each pixel, enabling the creation of a 3-dimensional data cube that integrates spatial and spectral information from the sample. While many spectrally variant fluorescent reporters are suitable for biological imaging, available fluorophores have broad excitation and emission spectra which makes distinguishing their individual contributions at every pixel a major challenge.

When there is overlap in the excitation and emission spectra of fluorophores, a single excitation wavelength band may excite more than one fluorophore and the emitted signals from different fluorophores can be recorded in the same emission bands. These phenomena is known as cross-talk and bleedthrough lead to inaccurate classification and quantification of signals from different fluorophores and thus hampers the ability to localize specific biological structures or molecules within cells and tissues. To overcome these issues, spectral imaging acquisition and analysis techniques, especially spectral unmixing methods, have been developed. These approaches aim to extract the spectral signatures of fluorophores from recorded images and determine the abundance of each fluorophore in every pixel. To tackle unmixing problems in different scenarios, various regularized learning methods have been developed in the literature (1–12). From a machine learning perspective, these methods essentially represent single-view learning where models are trained and predictions are made based on a single group of features that describes the field of interest, i.e., the emitted spectral profile of the fluorophores. However, organic fluorophores have not only unique emission spectral signatures, but also possess unique excitation spectral profiles. By recording the emission spectra with multiple combinations of excitation wavelengths in the same field, one can obtain multi-view data, each view of which can be considered as a distinct feature group of the field. In this study, we propose to address the biological spectral unmixing problem by developing a framework for a multi-view machine learning approach to biological spectral unmixing.

In the machine learning literature, it is observed that leveraging the complementary information available in different data views can help build more robust, flexible, and complex learning models with improved generalization ability (13–17). Such an observation gives birth to the introduction and development of multi-view machine learning approaches (18–20), which have demonstrated successful applications in various real-world scenarios (21–26). Motivated by the compelling empirical successes of multi-view learning in various application domains, we develop in this paper a framework for multi-view learning in the context of biological spectral unmixing by leveraging complete emission spectra obtained with various combinations of excitation wavelengths as a rich source of information on fluorophore distribution. The purpose is to significantly enhance the ability to discriminate fluorophores with highly overlapping spectra while allowing for a substantial expansion in the number of different fluorophores that can be employed and discriminated in a single experiment as their broad spectra lead to crowding and extensive bleed-through in the limited visible wavelength range. Within the context of biological spectral unmixing, there are some studies that make use of both excitation spectra and emission spectra. For instance, the study conducted in (27) demonstrated that using emission spectra at multiple excitation wavelengths can help distinguish 16 different fluorophores. By assuming that the acquired spectral image admits a three-way rank-one tensor representation, (7) proposed a blind source separation approach, which is essentially an unsupervised learning method, to address unmixing problems. Put differently, this approach relies on the assumption that emission spectra across different excitation wavelengths are scaled variations of each other. However, fluorophores absorb light energy at specific wavelengths and only emit light at longer wavelengths (28). As a result, signals with emission wavelengths shorter than an excitation wavelength are excluded, which may lead to insufficient exploration of the multi-view data. (29) used the integration of the emission spectra with different excitation wavelengths in unmixing. (30) investigated the biological spectral unmixing problem by specifically focusing on spectral image data acquired at two excitation wavelengths while assuming that the fluorophores of interest can not be correlated.

In contrast to previously reported work, we propose to make full use of the acquired multi-view image data to train learning machines. In particular, our proposed framework for multi-view learning in biological spectral unmixing is essentially a form of regularized learning that allows us to incorporate various noise, spatial, and sparsity constraints or other types of prior information about the fluorescent image. To validate the effectiveness of our proposed approach, we conducted experiments using simulated spectral images and real images of a microbial mixtures. The empirical findings unequivocally establish the superior performance of our framework over single-view learning, as demonstrated through both quantitative and qualitative analyses.

## Materials and methods

### Sample preparation

E. coli K12 (ATCC 10798) cells were grown to mid-log phase in Luria-Bertani LB Broth (Difco Laboratories, Inc.). E. coli cultures were fixed in 2% paraformaldehyde (EMS Diasum) at room temperature, then stored in 50% ethanol for at least 24 hours before FISH labeling as previously described (31). E. coli cells were labeled with the general bacteria probe, EUB338 (GCTGCCTCCCGTAGGAGT) conjugated to a fluorescent dye at the 5’ end.

### Imaging

Spectral images were acquired on a Zeiss LSM 980 confocal microscope with 32 anode spectral detector and a 63x 1.4 NA objective. Single-view images for comparison were acquired with a single combination of 488 nm, 561 nm, and 639 nm laser excitation wavelengths and multi-pass main beam splitter. The images were collected on the 32-anode spectral detector with 9.8 nm width spectral resolution in each channel. Multi-view images were acquired separately in the descending order of the excitation laser light wavelengths: 639 nm, 594 nm, 561 nm, 514 nm, 488 nm, and 445 nm. The number of channels with different excitation wavelengths or views are listed in Table 1.

**Table 1.** The number of channels with different excitation wavelengths. 488/561/639 represents the image recorded with a combination of 488 nm, 561 nm, and 639 nm laser excitation wavelengths.

| Laser (nm) | 445 | 488 | 514 | 561 | 594 | 639 | 488/561/639 |
| --- | --- | --- | --- | --- | --- | --- | --- |
| Channels | 28 | 24 | 21 | 15 | 12 | 7 | 24 |

Images were captured in a descending order of excitation laser light wavelengths, a strategic approach aimed at minimizing fluorophore bleaching (27). Because we record both the reference spectra and the unknown samples in the same sequence, moving from the longest to the shortest excitation wavelength, the impact of bleaching artifacts is minimized, as both the reference spectra and the unknown samples exhibit similar bleaching dynamics.

To minimize artifacts imposed by acquiring images with different main beam splitters, we align the images into one coordinate system using an intensity-based image registration algorithm (32). This algorithm optimizes the similarity between the images of the same field of interest through a 2D geometric transformation.

### Multi-view learning for biological spectral unmixing

A common assumption in spectral unmixing is that the signals recorded from various fluorophores within a pixel combine linearly. Linear spectral unmixing separates each pixel into its spectral signatures, referred to as endmembers, and their associated abundances.

Consider multi-view spectral images captured from the same field of interest, denoted as 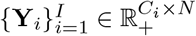. These im-ages comprise *C*_*i*_ channels and *N* pixels, obtained at *I* different combinations of excitation wavelengths. Let 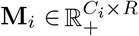 represent the endmember matrix associated with *R* fluorophores for the *i*-th combination of excitation wavelengths or the *i*-th view, and 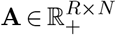 be the corresponding abundance matrix.

To accommodate such multi-view data in a linear unmixing context, we propose the following Multi-View Linear Mixture Model (MV-LMM):

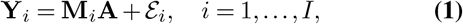

where 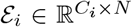 represents the unknown noise matrix from the *i*-th view. It reduces to the commonly used linear mixture model when *I* = 1, which corresponds to the scenario where only single-view data is available.

As is a common strategy in biological spectral imaging, we assume the availability of reference samples, each of which consists of a single fluorophore. We propose a two-step process of multi-view spectral unmixing, where the first step is to extract the endmember of each fluorophore from the reference spectral images recorded at different views, and the second step is, by using the extracted endmembers, to learn the abundances in a multi-view spectral image. A mathematical description of our proposed two-step approach can be detailed as follows:

### Step 1: Multi-view learning for endmember extraction

Denoting the multi-view reference images of a fluorophore recorded at different views as 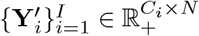, the endmember can be extracted through a multi-view machine learning scheme:

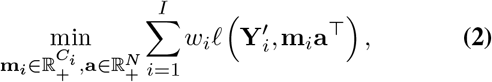

where **m**_*i*_ represents the endmember at the *i*-th view, **a** is the corresponding abundance vector, 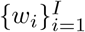 are the weights of different views, and 𝓁(·, ·) is a loss function.

### Step 2: Multi-view learning for abundance estimation

Recall that the endmember matrix **M**_*i*_ consists all fluorophores at the *i*-th view. Based on 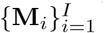 obtained from Step 1, in this step, we learn the abundances of the fluorophores in a given multi-view image set 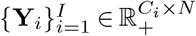 recorded at *I* views from the same field of interest. To this end, we propose the following multi-view machine learning method for abundance estimation:

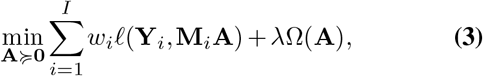

where *λ* is a tuning parameter and Ω(**A**) represents a penalty term imposed on **A**. Note that the weights of different views 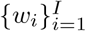 and the loss function 𝓁 in Eq. (3) can be different from those for the endmember extraction method Eq. (2).

This framework of multi-view machine learning effectively accommodates diverse prior information. For instance, the weights 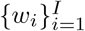 can be determined based on the varying importance of different views within specific applications. The choice of the loss function depends on the characteristics of the data. It can be selected as Poisson loss by assuming the presence of Poisson noise (7, 10), expectile loss (33) when dealing with asymmetric noise distribution, or other robust losses (34–36) for highly noisy data. The penalty term offers flexibility and can be chosen as the 𝓁_1_ norm and its variants to promote sparsity, nuclear norm to limit rankness, total variation for smoothing neighboring pixels, or a combination of several penalties. The parameter *λ* serves as the balancing factor, regulating the trade-off between the fidelity term and the penalty term.

## Experiments and results

We evaluated the effectiveness of our proposed multi-view learning by comparing it with a commonly used single-view learning approach, utilizing the combination of 488 nm, 561 nm, and 639 nm laser excitation wavelengths. To maintain fairness and simplicity in the comparison, we set the weights in Eq. (2) and Eq. (3) as equal and opt for the least squares loss function with no penalty term. This choice eliminated the need for a tuning process. Then Eq. (2) is expressed as

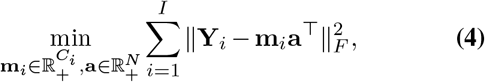

where ‖ · ‖ represents the Frobenius norm. And Eq. (3) is formulated as

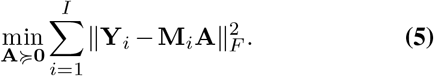

Both optimization problems Eq. (4) and Eq. (5) were addressed using an Alternating Direction Method of Multipliers (ADMM) technique (37) to ensure efficient and effective solutions. For single-view learning, we employed the Nonnegative Matrix Factorization (38) algorithm for endmember extraction and the Nonnegative Least Squares (NLS) unmixing method for abundance estimation (39).

### Simulations of 100 endmembers

To assess the performance of our proposed methods across varying numbers of views in the presence of numerous highly overlapping endmembers, we generated 100 endmembers using a Gaussian distribution with 24 channels, as depicted in Figure 1. The correlation coefficient between any two adjacent endmembers is 0.973, indicating significant spectral overlap. With these endmembers, we simulated 1000 spectral images, each comprising 1000 pixels. Within each pixel, the abundance of one random endmember was generated from a uniform distribution U(0, 1), while the abundances of the remaining endmembers were set to zero. Additionally, the endmembers were modified by randomly generated excitation intensities, which follow a uniform distribution U[0, 1]. To simulate realworld conditions, Poisson noise with a signal-to-noise ratio (SNR) of 5 and Gaussian noise with an SNR of 40 were introduced into the generated data.

**Fig. 1.**
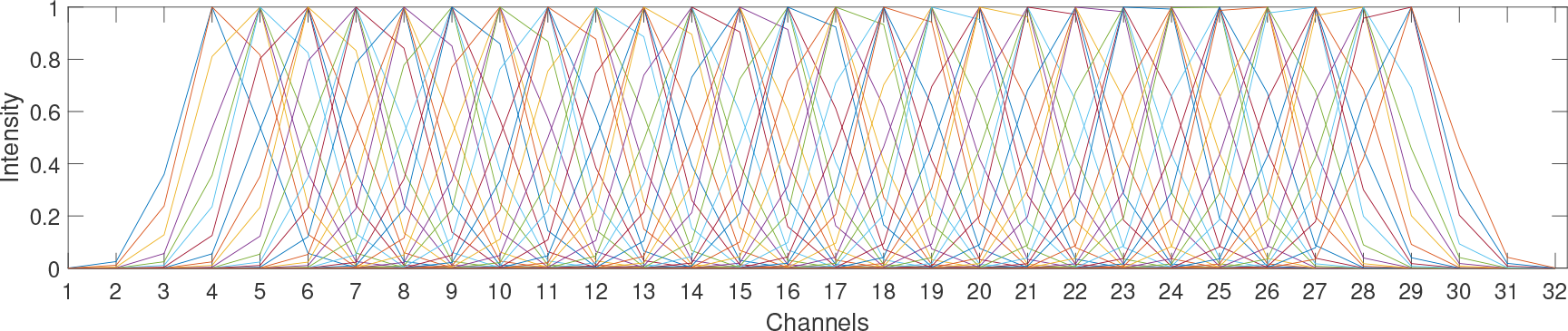
100 endmembers with 32 channels were created through a Gaussian distribution characterized by close means and a standard deviation of 0.5. Each color represents a distinct endmember.

To evaluate the accuracy of unmixing with *N* pixels and *R* endmembers, we employed the Root Mean Square Error (RMSE) criterion, defined as follows:

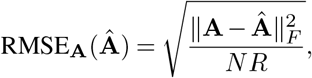

where **A** represents the simulated abundance matrix, which serves as the ground truth, and Âdenotes the estimated abundance matrix. The RMSE criterion measures the dissimilarity between the true abundance matrix and the estimated abundance matrix. Figure 2 presents the boxplots of RMSEs obtained from simulations with varying numbers of views. The RMSEs exhibit a decreasing trend as the number of views increases. The average RMSEs show a significantly lower value with a higher number of views compared to cases with a lower number of views.

**Fig. 2.**
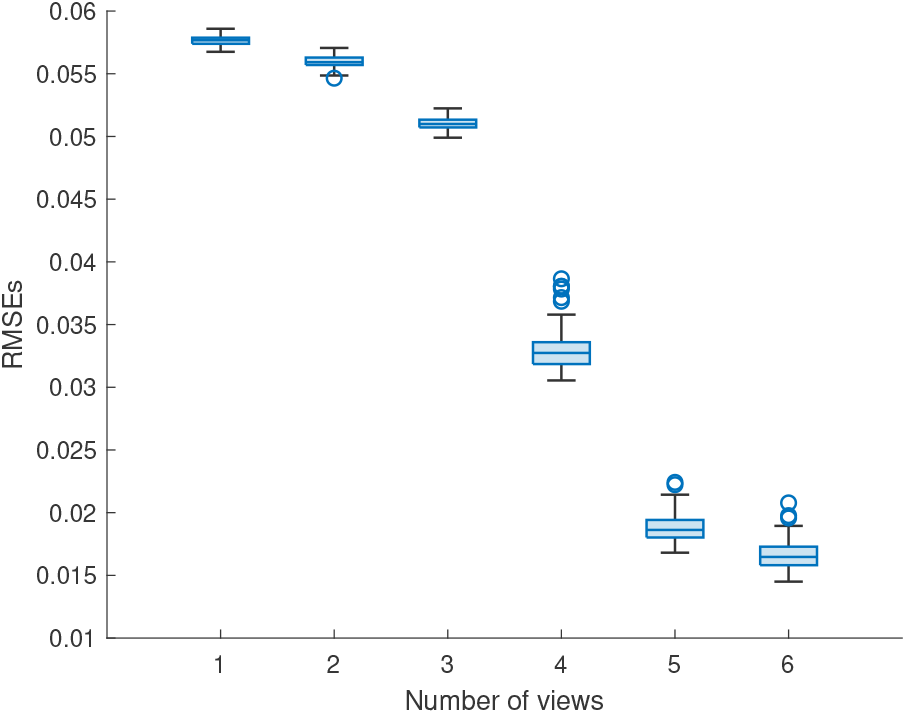
Boxplots of Root Mean Square Errors (RMSEs) were derived from simulations involving varying numbers of views. Each RMSE was assessed based on the unmixing results obtained from simulated spectral images, where the number of views corresponds to the quantity of simulated spectral images.

### Real biological images

In this section, an assessment of multi-view learning performance was conducted on real biological images of fluorescently labeled E. coli cells. E. coli cells were labeled in a fluorescence in situ hybridization (FISH) procedure with the general bacterial probe. Twelve versions of the same oligonucleotide FISH probe were synthesized, each version conjugated to a different fluorophore endmember, characterized by significant overlapping excitation and emission spectra, as illustrated in Figure 3. Endmember spectral profiles were extracted from their corresponding multi-view reference images through Eq. (4) from reference images of pure populations of the E. coli, each population labeled with only one version of the FISH probe. Then the abundances of these fluorophores were reconstructed from an additional set of reference images. It was assumed, during the unmixing process, that all endmembers were present in each image, despite only a single endmember existing in each reference image.

**Fig. 3.**
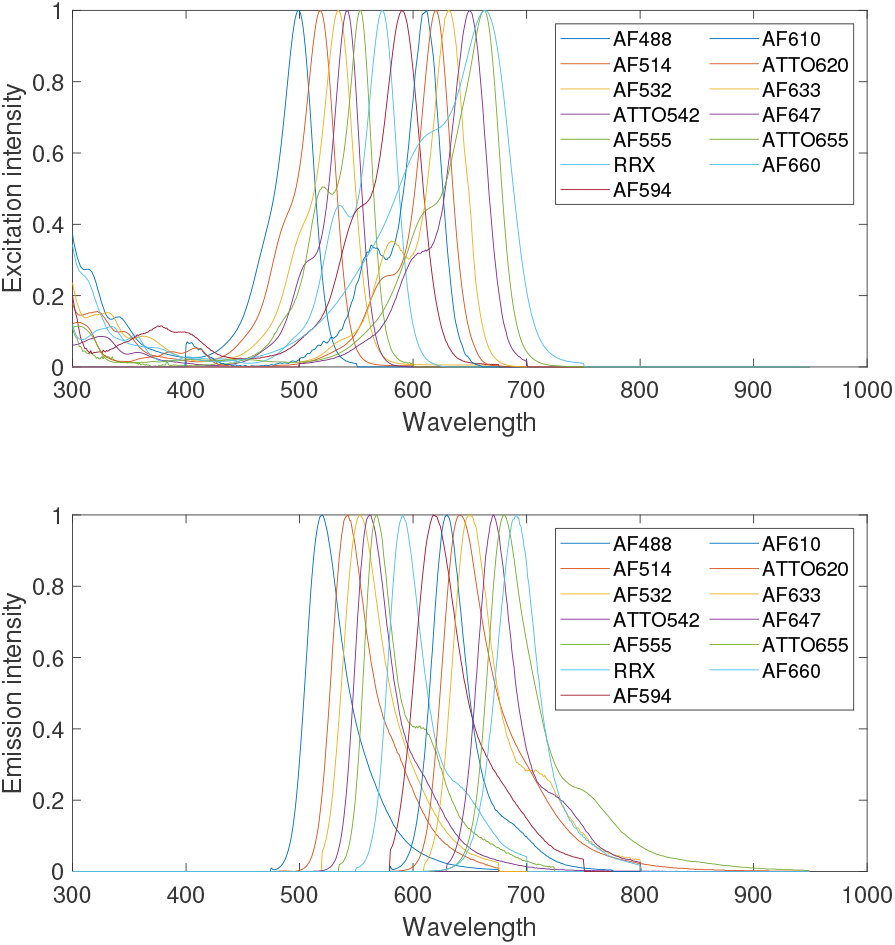
Standardized overlapping (top) excitation and (bottom) emission spectra of 12 Fluorophores, with AF representing Alexa Fluor and RRX denoting Rhodamine Red-X.

To evaluate the unmixing results, a pixel in a reference image is correctly identified if the estimated abundance of the corresponding fluorophore is more than that of all other fluorophores. To visually represent the performance of the unmixing results, each of the twelve fluorophores was assigned a distinct color. For every pixel, the color of the fluorophore with the highest calculated abundance among all fluorophores was assigned to that pixel. This color assignment scheme facilitated a clear and intuitive depiction of the dominant fluorophore in each pixel, thereby providing a visual representation of the unmixing results. The estimated abundances of the twelve reference images, obtained through our proposed multi-view learning and the single-view learning, are presented in Figure 4, respectively. As depicted in the figures, our proposed method exhibits superior accuracy, with a greater number of correctly identified pixels compared to the single-view learning. Notably, in Figure 4(c), the multiview learning for AF532 demonstrates a significantly better outcome than the corresponding single-view learning. This improvement is attributed to the substantial excitation intensity of AF532 at wavelengths 488 nm, 514 nm, and 532 nm. Despite the reduction in number of channels with increasing wavelength, the rich information from multi-view learning aids in effectively distinguishing AF532 from other highly overlapping fluorophores. Despite the utilization of six excitation wavelengths respectively, the number of views for each fluorophore is not as extensive due to the limited range of nonzero excitation spectra and the reduction in number of channels. Consequently, in Figure 4(k), multi-view learning for the very long wavelength ATTO655 yields a result similar to that of single-view learning.

**Fig. 4.**
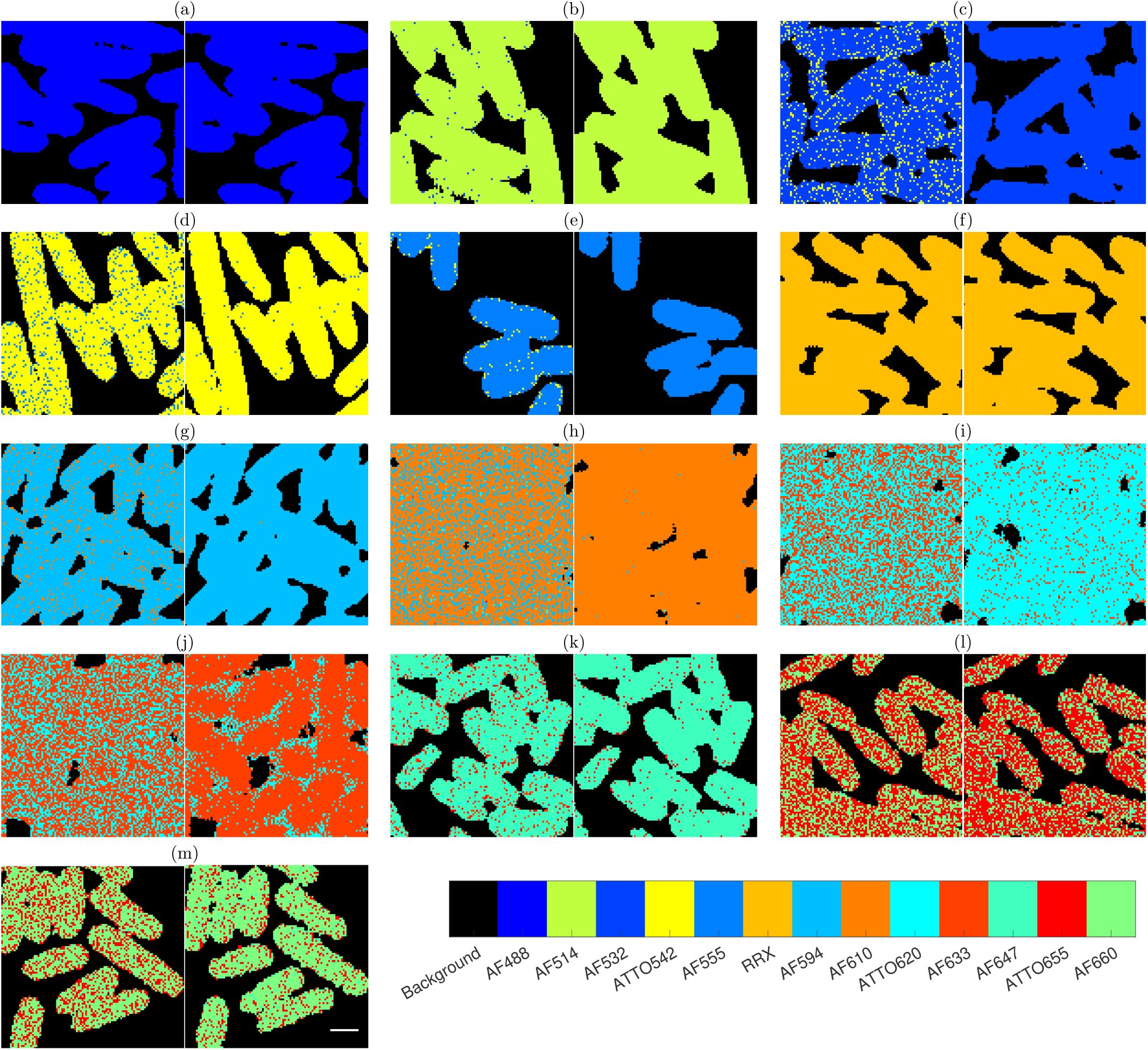
Abundances of reference images of (a) AF488, (b) AF514, (c) AF532, (d) ATTO542, (e) AF555, (f) RRX, (g) AF594 (h) AF610, (i) ATTO620, (j) AF633, (k) AF647, (l) ATTO655, and (m) AF660 estimated by (left of all pairs) single-view learning and (right of all pairs) multi-view learning. The white rectangular bar in the lower right corner of the right image of (m) represents a scale of 1 *µ*m. Each pixel’s color corresponds to the fluorophore with the highest estimated abundance, as indicated by the legend with AF representing Alexa Fluor and RRX denoting Rhodamine Red-X.

We proceeded to reconstruct the abundances of the fluorophores in a mixture sample of E. coli cells, characterized by similar morphology as the reference samples. The estimated abundances obtained through multi-view learning and single-view learning are visualized in Figure 5. Notably, the abundance matrix derived from multi-view learning reveals distinct oval shapes, each corresponding to the same color or the highest abundance of a specific fluorophore. This contrasts with the abundance matrix obtained through singleview learning.

**Fig. 5.**
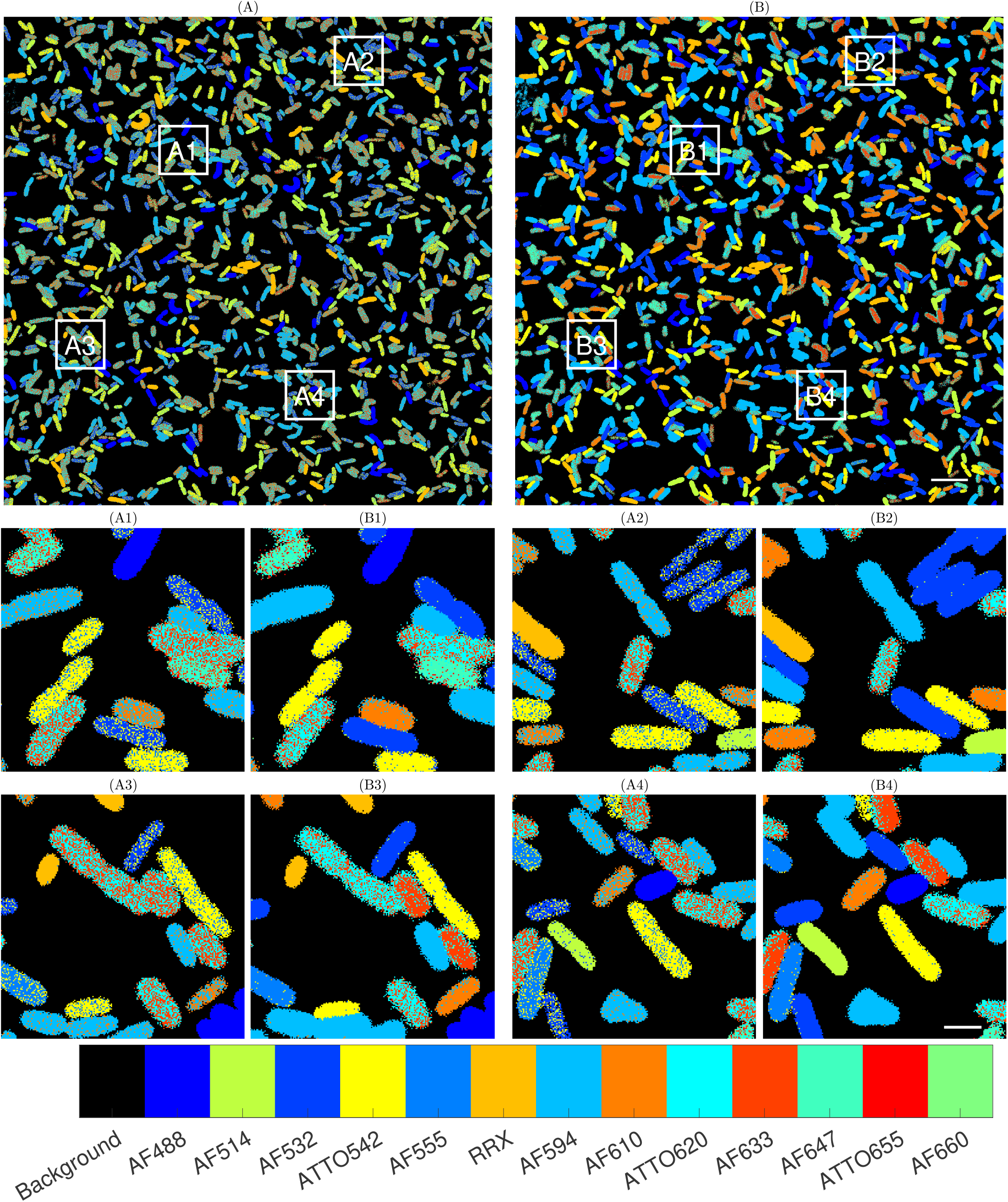
Abundances of mixed E. coli spectral image estimated by (A) single-view learning and (B) multi-view learning. (A1)–(A4) are magnified views of the regions within the four white boxes in (A) and (B1)–(B4) are magnified views of the regions within the four white boxes in (B). The white rectangular bar in the lower right corner of (B4) represents a scale of 2 *µ*m. Each pixel’s color corresponds to the fluorophore with the highest estimated abundance, as indicated by the legend with AF representing Alexa Fluor and RRX denoting Rhodamine Red-X.

## Conclusion

We designed a multi-view machine learning framework to effectively differentiate fluorophores with significant spectral overlap. The multi-view data was acquired by recording the emission spectra with various combinations of excitation wavelengths. Through our proposed multi-view machine learning approach, one can perform endmember extraction and abundance estimation using Eq. (2) and Eq. (3).

Our approach demonstrated exceptional accuracy in estimating fluorophore abundances, particularly for those with highly overlapping emission spectra, through the application of multi-view learning. Simulated data results highlighted that incorporating more views led to more accurate unmixing outcomes. The unmixed real biological spectral images further underscored the efficacy of our proposed approach. Note that the simulation utilized only 100 endmembers as an example, and the approach can handle a variable number of endmembers as required.

It is important to note that we opted for least squares unmixing methods Eq. (4) and Eq. (5) for their simplicity and computational efficiency. However, other data fidelity terms tailored to the specific dataset could be considered. Additionally, the constraints in Eq. (3) on abundances may be applied based on the particular requirements of the application. As a first approach, our multi-view framework incorporates multiple discrete wavelength excitations as different views of the data, because it is well appreciated that organic fluorophores have unique excitation and emission spectra. As a framework the approach presented here could, in future, incorporate other types of data as different views of the endmembers including fluorescence lifetime information and other characteristics that are unique to each endmember used in any single biological fluorescence imaging experiment.

## Author contributions statement

Y.F. and A.M.V. conceived the method. R.W. conducted the experiments, analysed the results, and wrote the manuscript. Y.F. and A.M.V. reviewed the manuscript.

## Supplementary Note 1: Funding

This work was supported by National Institutes of Health [grant numbers R01DE031213, R01DE030927, S10OD028600]; and National Science Foundation [grant numbers 1636933, 2111080, 1920920].

